# Requirement of GrgA for *Chlamydia* infectious progeny production

**DOI:** 10.1101/2023.03.09.531986

**Authors:** Bin Lu, Yuxuan Wang, Wurihan Wurihan, Sydney Yeung, Joseph D. Fondell, Xiang Wu, Huizhou Fan

## Abstract

Hallmarks of the developmental cycle of the obligate intracellular pathogenic bacterium *Chlamydia* are the primary differentiation of the infectious elementary body (EB) cell type into the proliferative reticulate body (RB) and the secondary differentiation of RBs back into EBs. The detailed mechanisms regulating these transitions are unclear. In this study, we developed a novel strategy termed DOPE (dependence on plasmid-mediated expression) that allows for the knockdown of essential genes in *Chlamydia*. Importantly, we demonstrate that GrgA, a *Chlamydia*-specific transcription factor, is essential for the secondary differentiation of RBs into EBs. Our development of a conditional GrgA-deficient chlamydiae should prove valuable for future studies examining chlamydial growth, development, and pathogenicity. Furthermore, because EB formation is absolutely required for chlamydial dissemination within infected individuals, and for chlamydial transmission to new hosts, our maturation-defective chlamydiae system may serve as an attractive attenuated vaccine methodology for the prevention of chlamydial diseases.

## INTRODUCTION

*Chlamydia* is an obligate intracellular bacterium with a unique developmental cycle ^1-3^. The chlamydial developmental cycle is characterized by two morphologically and functionally distinct cell types termed the elementary body (EB) and reticulate body (RB). With limited metabolic activity, the small, electron-dense EB is capable of temporarily surviving extracellular environments and is responsible for infecting host cells to initiate the developmental cycle. After entering host cells, Ebs increase in size and differentiate into dividing RBs. After several rounds of replication, RBs asynchronously differentiate back into non-replicating EBs ^3,4^. After release from host cells, progeny EBs either infect additional cells within the same host or are transmitted to new hosts. Unlike EBs, any RBs released from host cells are unable to initiate new developmental cycles but can induce immune responses upon decomposition inside host cells.

The chlamydial developmental cycle is transcriptionally regulated ^3,5,6^. After EBs enter host cells, early genes are activated during the first few hours enabling primary differentiation into RBs. Starting at around 8 hours post-infection, midcycle genes, representing the vast majority of all chlamydial genes, are expressed enabling RB replication. At around 24 hours post-infection, late genes are activated to enable the secondary differentiation of RBs back into EBs.

As a subunit of the RNA polymerase (RNAP) holoenzyme, sigma factor recognizes and binds specific DNA gene promoter elements allowing RNAP to initiate transcription ^7-9^. *Chlamydia* encodes three sigma factors termed σ^66^, σ^28^, and σ^54 10-12^. σ^66^ RNAP holoenzyme is involved in the expression of early, mid, and late genes, whereas the σ^28^ RNAP and σ^54^ RNAP are responsible for transcribing only a subset of late or mid-late genes ^13-15^. A small number of genes have tandem promoters, and their expression are regulated by multiple sigma factors ^13^.

GrgA is a *Chlamydia-*specific transcriptional regulator identified through promoter DNA pulldown that binds both σ^66^ and σ^28^ and activates the transcription of multiple chlamydial genes *in vitro* and *in vivo* ^16-18^. RNA-Seq analysis of *C. trachomatis* conditionally overexpressing GrgA, together with GrgA *in vitro* transcription assays, have identified two other transcription factor-encoding genes, *euo* and *hrcA*, as members of the GrgA regulon ^16^. Both immediate-early genes, *euo* and *hrcA* are transcribed during the early phase and midcycle ^16^. While Euo is a repressor of chlamydial late genes ^19^, HrcA regulates the expression of multiple protein chaperones ^20^, which are essential for bacterial growth ^21^. These findings suggest that GrgA regulates early chlamydial development.

To further determine the role of GrgA in chlamydial physiology, we attempted but failed to disrupt *grgA* through group II intron (Targetron ^22^) insertional mutagenesis. Previously, saturated chemical mutagenesis also failed to generate *grgA-*null mutants ^23^. Given that Targetron and chemical mutagenesis have been successfully used to disrupt numerous non-essential chlamydial genes ^23-28^, our finding suggest that *grgA* is an essential gene. In this work, we confirm that *grgA* is indeed an essential gene by developing and applying a novel genetic tool that we term DOPE (dependence on plasmid-mediated expression). Importantly, we show that GrgA is necessary for RB-to-EB differentiation during the late developmental cycle and is further required for optimal RB growth. To the best of our knowledge, this is the first report implicating the requirement of a single chlamydial regulatory factor in the formation of progeny EBs.

## RESULTS AND DISCUSSION

### DOPE (dependence on plasmid-mediated expression) enables *grgA* disruption

Targetron, a group II intron-based insertional mutagenesis technology, has been used to successfully disrupt numerous chlamydial chromosomal genes ^24-28^. In an effort to knock out GrgA expression in *Chlamydia* and investigate its physiological actions, we utilized Targetron vectors containing spectinomycin-resistance gene-bearing group II introns specific for multiple *grgA* insertion sites. Despite several attempts, we failed to generate any *grgA*-null mutants. Indeed, spectinomycin-resistant chlamydiae were obtained from only two transformed cultures, yet diagnostic PCR analysis failed to demonstrate insertion of the group II intron into *grgA* indicating nonspecific targeting. Together with the failure to obtain *grgA-*null mutants by the Valdivia Lab, our failure to establish stable *grgA*-null mutants using Targetron insertional mutagenesis suggested to us that *grgA* may be essential for chlamydial growth and development.

As a new and alternative approach to investigate the biological functions and underlying mechanisms of *grgA* and other genes essential for chlamydial growth and/or development, we developed the dependence on plasmid-mediated expression (DOPE) tool (Fig.1). Because essential genes are required for normal chlamydial growth or development, their disruption in wildtype chlamydiae will result in lethality (Fig. 1 A, B). However, transformation of wildtype *Chlamydia* with a recombinant plasmid carrying the essential chromosomal gene downstream of an inducible promoter allows for the disruption of the chromosomal allele when the inducer is present in the culture medium (Fig. 1C). In the resulting essential gene-disrupted strain, withdrawal of the inducer will cause depletion of the gene products generated from the recombinant plasmid and allow for functional and mechanistic analyses of the essential gene (Fig. 1D).

**Fig. 1.**
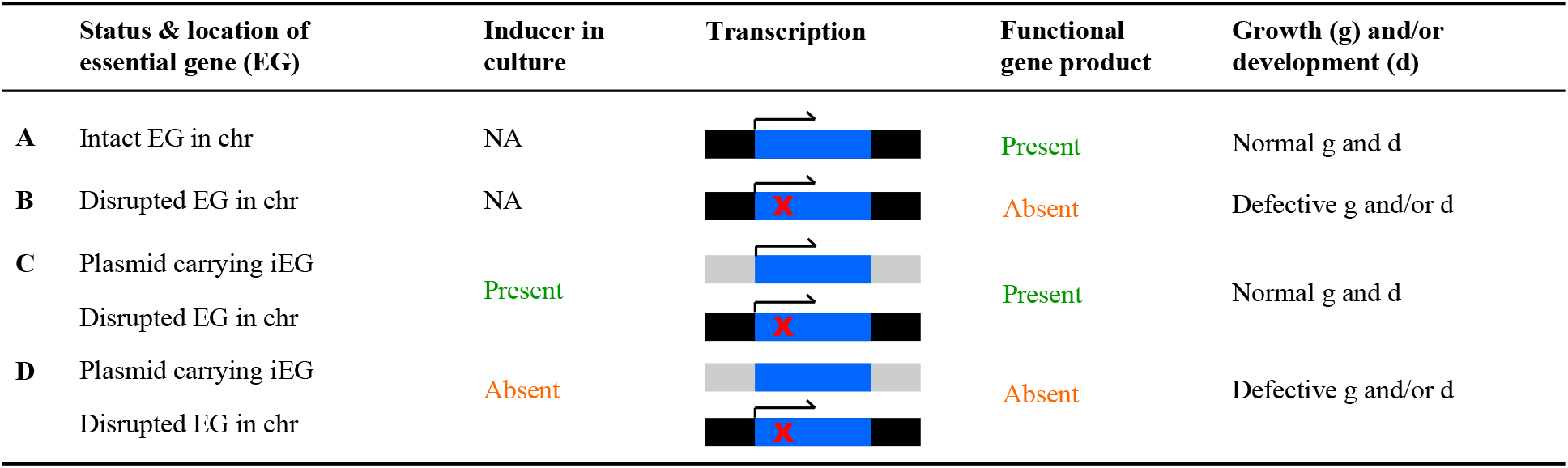
The DOPE (Dependence On Plasmid-mediated Expressed) technology allows for functional and mechanistic interrogation of chromosome-encoded essential genes. Dark and grey lines signify chromosomal and plasmid sequences, respectively. Arrows indicate transcription. Abbreviations: EG, essential genes; iEG, inducible EG; chr, chromosome; NA, not applicable; g, growth; d, development.

To apply DOPE to the study of *grgA*, we first constructed a plasmid encoding an anhydrotetracycline (ATC)-inducible *grgA* allele (plasmid-encoded inducible *grgA* or peig) (Fig. S1). Compared to the native chromosomal *grgA* allele that contains a group II intron-target site between nucleotides 67 and 68, the *grgA* allele in peig carried a His-tag sequence and four synonymous point mutations surrounding the group II intron-targeting site (Fig. 2A). We transformed wildtype *C. trachomatis* with an intact chromosomal *grgA* (L2/cg) with peig to derive L2/cg-peig. We next transformed L2/cg-peig with the aforementioned Targetron plasmid carrying an *aadA-*containing group II intron with the insertion site between nucleotides 67 and 68 in *grgA* (Fig. S2). Since the Targeton target site in the peig allele has been mutated, the vector can only insert into the chromosomal *grgA* allele (Fig. 2A). Following transformation of L2/cg-peig with the Targetron plasmid, we supplemented the culture medium with ATC and spectinomycin to induce GrgA expression from peig while selecting for chromosomal mutants carrying an intron-disrupted *grgA*. PCR analysis confirmed the chlamydial genotypes L2/cg, L2/cg-peig, as well as the plasmid-complemented, chromosomal *grgA-*disrupted L2/cgad-peig (Fig. 2A, 2B and Table S1). Tracings of Sanger sequences in Fig. 2C confirmed the nucleotide sequences surrounding the intron-target site in L2/cg and L2/cg-peig and the *grgA-*intron joint regions in L2/cgad-peig. As expected, western blotting detected ATC-independent chromosome-encoded GrgA expression and ATC-dependent plasmid-encoded His-GrgA expression in L2/cg-peig (Fig. 2D). Significantly,western blotting also detected time-dependent GrgA loss in L2/cgad-peig upon ATC withdrawal (Fig. 2E). By 2 h post-ATC withdrawal, His-GrgA became nearly undetectable. Taken together, the data presented in Fig. 2B-E demonstrate that we successfully disrupted wildtype chromosomal *grgA* in L2/cgad-peig, in which expression, and thus function, of GrgA depends on ATC-induced GrgA expression from the peig plasmid.

**Fig. 2.**
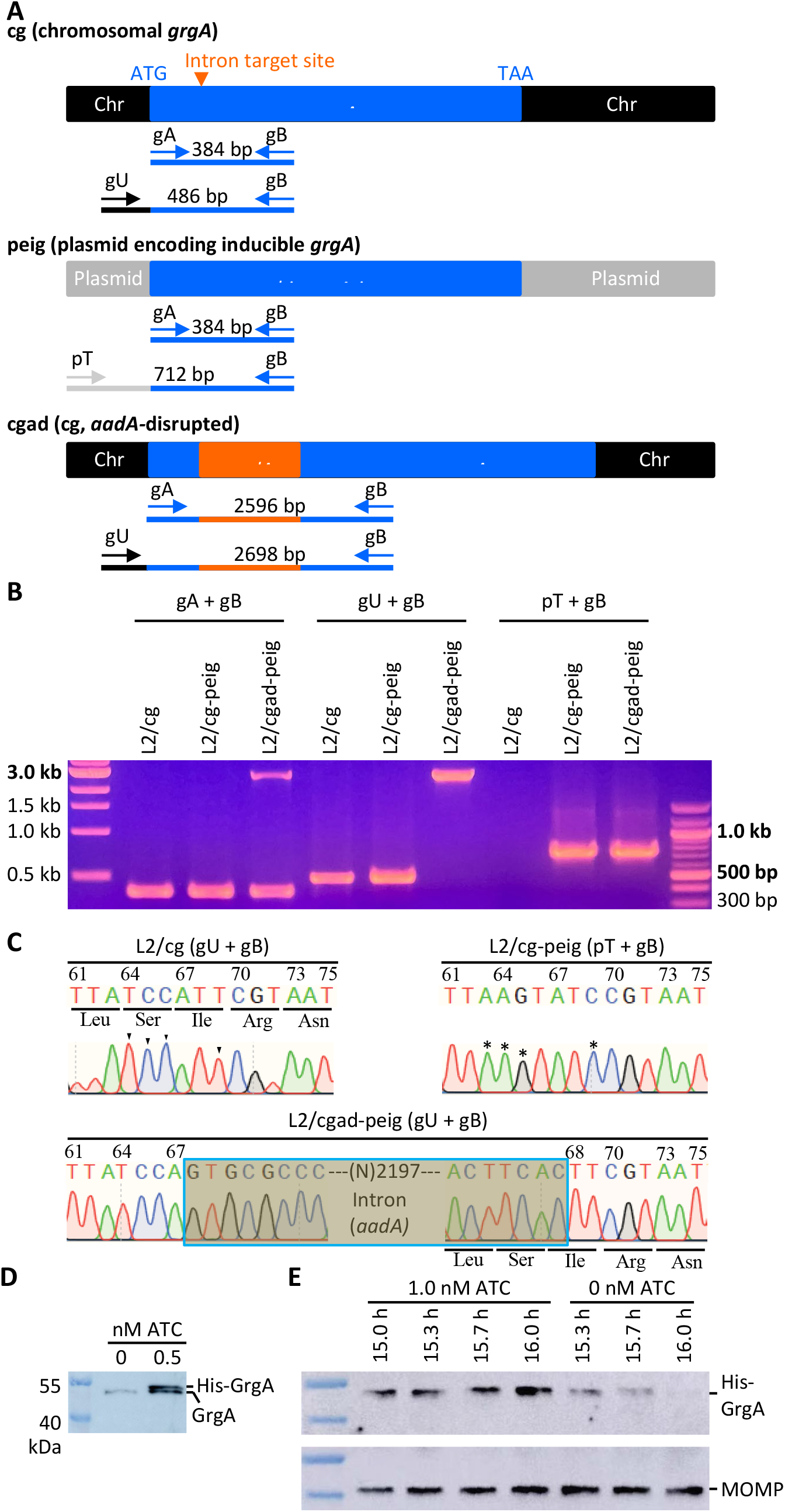
Confirmation of the disruption of the chromosome-encoded *grgA* by group II-intronand re-expression of GrgA from a transformed plasmid in the DOPE system. (A) Schematic drawings of *grgA* alleles, locations of intron-target site,diagnostic primers, and size of PCR products obtained with different sets of primers. Abbreviations: itsm, intron target site mutated; Chr, chromosome. (B) Gel image of PCR products amplified with DNA of wildtype *C. trachomatis* L2 (L2/cg), L2/cg transformed with the *his-grgA-itsm* expression plasmid (L2/cg-peig), and L2 with *aadA-*300 bp disrupted chromosomal *grgA* complemented with peig (L2/cgad-peig) using primer sets shown in (A). (C) Sanger sequencing tracings of PCR products showing the intron-target site in L2/cg, mutations surrounding this site conferring resistance to intron targeting in peig, and *grgA-*intron joint regions in the chromosome of L2/cg-peig. Wildtye bases and corresponding mutated bases are shown with arrowheads and asterisks, respectively. (D) Western blotting detection only chromosome-encoded GrgA in L2/cg-peig cultured in ATC-free medium and both chromosome-(*aadA)* encoded GrgA and plasmid-encoded His-GrgA in L2/cg-peig cultured in the ATC-containing medium for 14 h. (E)Western blotting showing time-dependent loss of His-GrgA in L2/cgad-peig upon ATC MOMP withdrawal. The membrane was first probed with an anti-major outer membrane protein, striped, and then reprobed with an anti-GrgA antibody.

### GrgA-deficient chlamydiae display slower growth and fail to form progeny EBs

To determine the effect of GrgA deficiency on *C. trachomatis* growth and development, we compared the growth of L2/cgad-peig cultured in media with and without 1 nM ATC. Interestingly, immunofluorescence using an antibody specific for the major outer membrane protein revealed a slower growth in ATC-free cultures as determined by the inclusion size (Fig. 3A). Quantitative PCR analysis further showed slower genome replication in the ATC-free cultures (Fig. 3B, Fig. S3). The genome doubling time of L2/cgad-peig in the ATC-containing cultures was 2 h (Fig. 3B), which is identical to the doubling time of wild L2 in our lab ^16^. In contrast, the genome doubling time of of L2/cgad-peig in ATC-free culture was 4 hours (Fig. 3B). Remarkably, ATC-free cultures exhibited a severe deficiency in the production of infectious progeny EBs (Fig. 3C). ATC-containing cultures produced on the order of >10^4^ EBs at 24 h, > 10^6^ at 34 h, and nearly 5 × 10^6^ at 48 and 72 h (Fig. 3C). By stark contrast, ATC-free cultures produced no detectable EBs at 24 h and only single digit EBs at 34 h and thereafter (Fig. 3C), even though the chlamydial genome copy number in the ATC-free cultures at 34 h slightly exceeded that in the ATC-containing cultures at 24 h (Fig. 3B). Consistent with the EB quantification assays, ultra-thin section transmission electron microscopy readily detected EBs at 36 h in the ATC-containing cultures and were predominant at 45 h (Fig. 4). In contrast, EBs were rarely found in ATC-free cultures at 35, 45, or 60 h (Fig. 4; Fig. S4). Taken together, these findings demonstrate that GrgA deficiency plays an important role in optimal RB growth and is crucial for the production of infectious EBs from RBs.

**Fig. 3.**
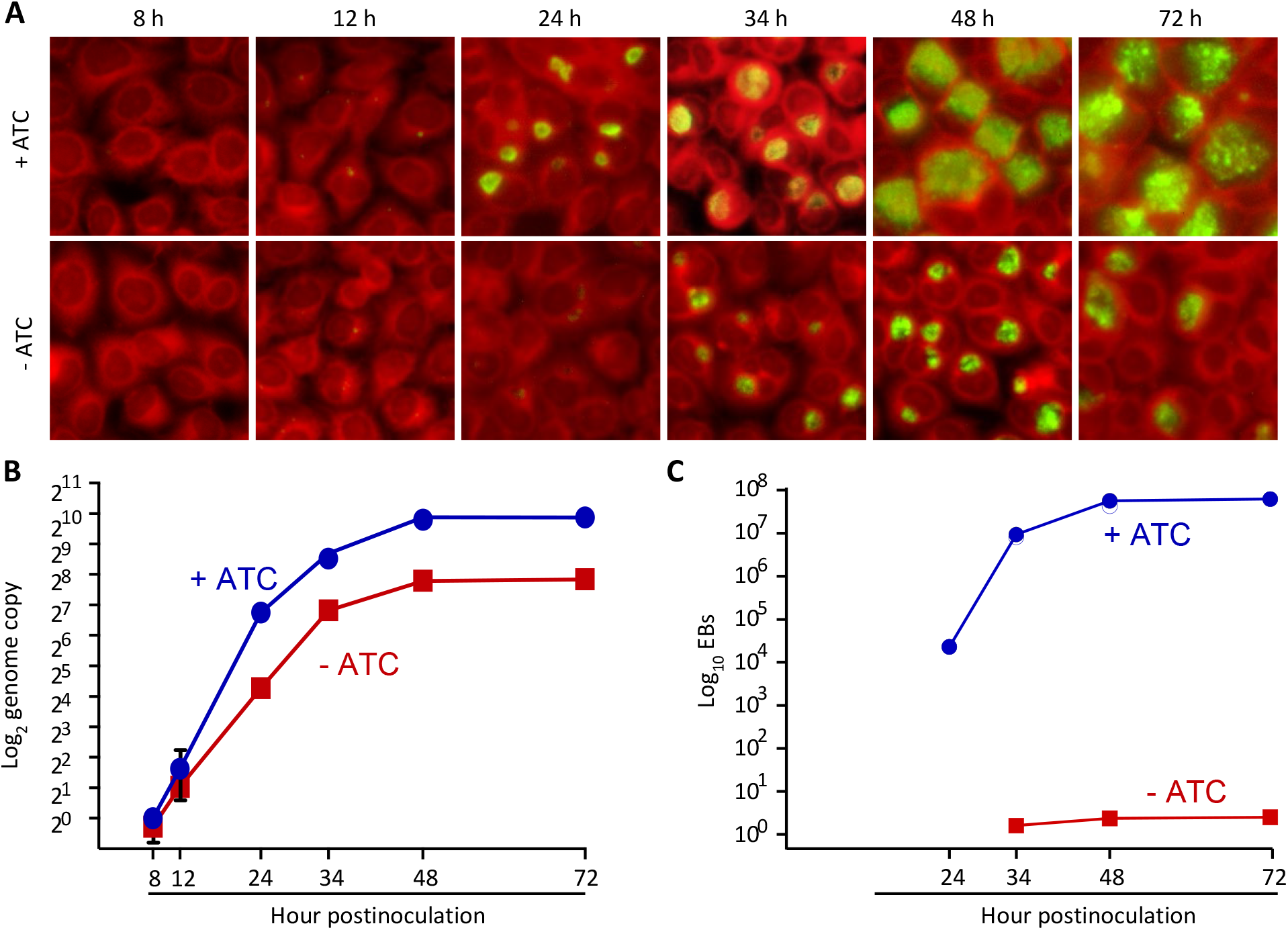
GrgA deficiency leads to slowed RB growth and lack of progeny EB formation. L2/cg-peig-infected HeLa cells were cultured in the presence or absence of 1 nM ATC. At indicated hpi, cultures were terminated for immunofluorescence assay (A), genome copy quantification (B), or EB quantification (C). (B, D) Data represent averages ± standard deviations of triplicate cultures.

**Fig. 4.**
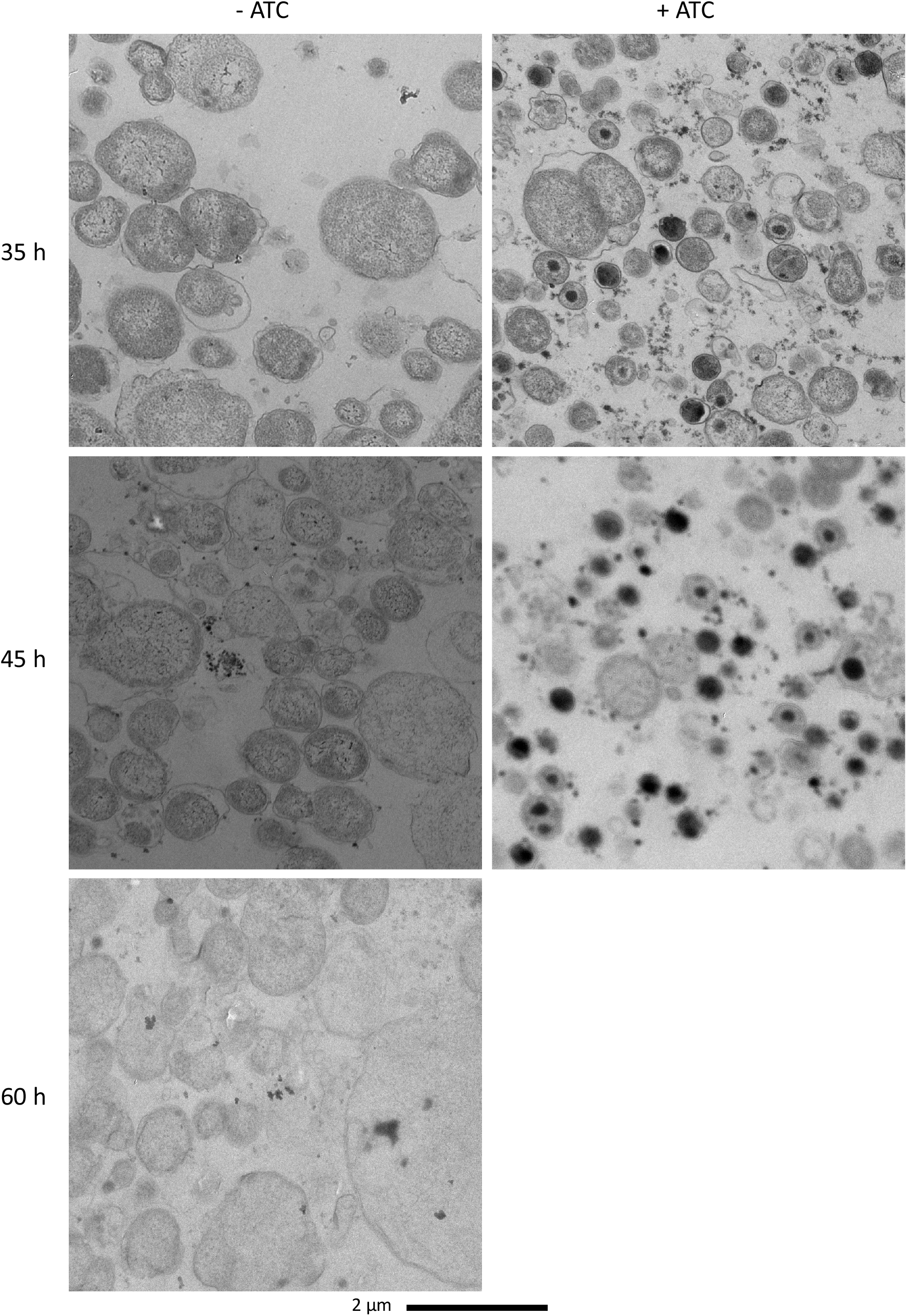
Electron microscopy demonstrates lack of EBs in L2/cgad-peig cultured in ATC-free medium at 35, 45, and 60 h. 1 nM ATC cultures at 60 h was not processed for EM because most inclusion already burst by that point. Note that EBs are ~400 nm diameter cellular forms with high electron density found in ATC-containing cultures. Small irregularly-shaped dark particles in both ATC-containing and ATC-free cultures are glycogen particles.

### *tetR* mutations enables GrgA expression and EBs to escape in the absence of ATC

Whereas GrgA-null chlamydiae display a severe deficiency in the formation of EBs, we were able to detect an extremely low trace background of EBs in chlamydial cultures lacking ATC (Fig. 3C). We hypothesized that mutations in the *tetR* gene and/or mutations in tetO (TetR operator) might disable the ability of TetR to repress *grgA* in L2/cgad-peig and consequently allow EBs to form in the absence of ATC. To investigate this theory, we recovered plasmids from EBs formed in ATC-free cultures and expanded them in *E. coli*. Notably, DNA sequencing showed that a single nucleotide polymorphism (SNP) in the *tetR* gene had occurred in all 10 plasmids. Three distinct SNPs were detected; their locations and effects on the 208-aa TetR protein are shown in Fig. 5. Two of the SNPs cause premature termination at codons 16 or 158, while the third causes a frame shift at codon 64. These findings indicate that lack of full-length TetR, or synthesis of a functionally defective TetR, results in the leaky expression of wildtype GrgA in the absence of ATC. Taken together with findings in Figs. 3 and 4, this data strengthens our conclusion that GrgA is required for EB formation.

**Fig. 5.**
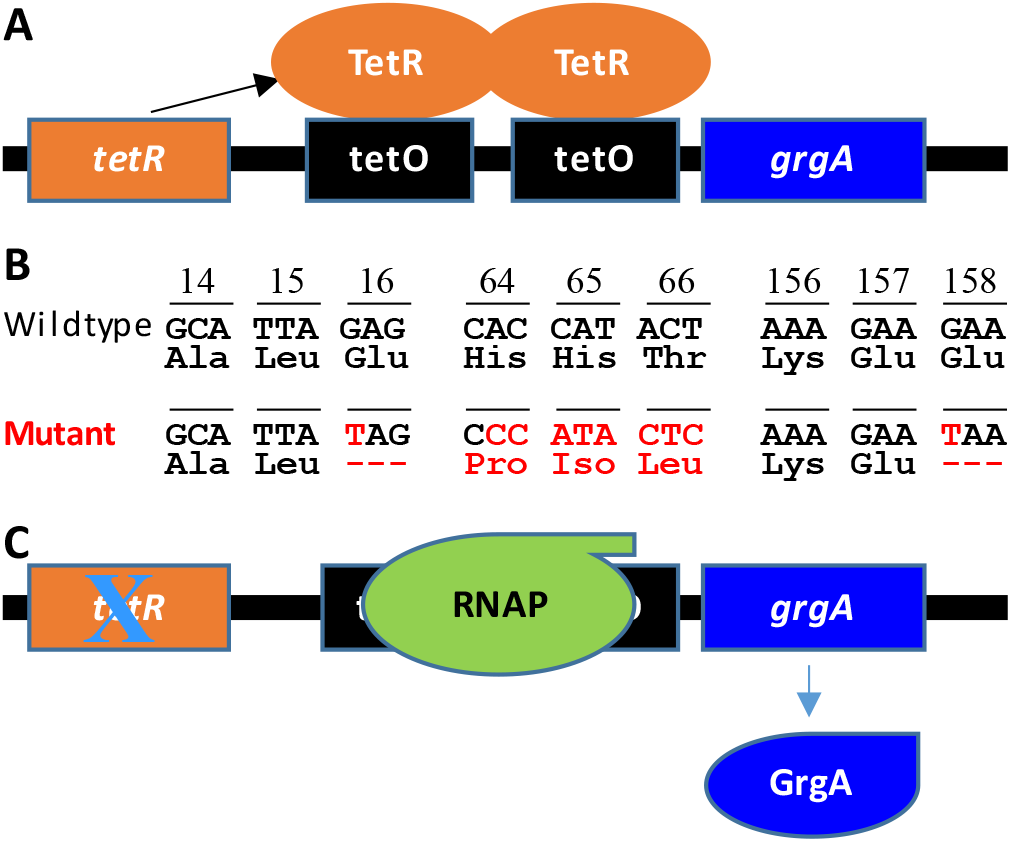
Failed *grgA* repression in peig is responsible for EB escape in ATC-free cultures of L2/cgad-peig. (A) In L2/cgad-peig, the plasmid allele of *grgA* is repressed in the presence of ATC. (B) Mutations identified in *tetR* in the plasmids isolated from L2/cg-peig EBs formed in the absence of ATC lead to premature translation termination or frameshift. Codon positions are indicated. Codon positions are indicated. Wild -type DNA and amino acids are shown in black and mutated or frame -shifted nucleotides as well as consequent translational effects in red. (C) Mutations detected in (B) causes a loss in *grgA* repression, which enables peig to express functional TetR and consequentEB formation.

## DISCUSSION

Since the first publication demonstrating reproducible transformation of *Chlamydia* with a shuttle vector 12 years ago ^29^, the *Chlamydia* research community has utilized the reverse genetic tool to investigate gene function through ectopic gene overexpression, gene knockdown, and other approaches^13,24-28,30,31^. Nonetheless, effective strategies for disrupting or depleting truly essential genes have hampered research in *Chlamydia* and other biological systems. In this report, we developed a novel tightly regulated inducible expression system termed DOPE that allows for the knockdown of essential genes in *Chlamydia*. The DOPE system not only represents a convenient and versatile tool for establishing the essentiality of genes, but also defining their underlying mechanisms. Unlike CRISPR interference, DOPE lacks off-target effects or general nonspecific toxicity ^32-34^.

The inability to generate *grgA*-null mutants by us using gene target mutagenesis and by the Valdivia Lab using random mutagenesis strongly indicated that *grgA* is crucial for *Chlamydia* growth and viability. Indeed, by applying DOPE strategy to knockdown GrgA expression, we show here that GrgA plays a critical role in maintaining RB replication efficiency and is absolutely essential for the RB-to-EB differentiation. Surprisingly, even though previous studies demonstrated that two immediate-early transcriptional factors *euo* and *hrcA* are readily upregulated following the induction of GrgA overexpression ^16^, genome replication kinetics data presented in Fig. 3B suggests that the primary EB-to-RB differentiation is not affected by ATC omission in the culture. However, these data do not exclude the possibility that GrgA plays a role in the primary differentiation because the amount of GrgA prepacked into EBs could be sufficient for supporting the primary EB-to-RB differentiation. In keeping with this view, both we and the Valdivia Lab detected significant amounts of GrgA in *C. trachomatis* EBs ^18,35^. Our ongoing transcriptomic analysis will elucidate the mechanisms by which GrgA regulates RB growth and RB-to-EB differentiation.

Formation of EBs is absolutely required for dissemination of chlamydial infection within the infected host and transmission to new hosts. Because RBs and EBs share most of the immunodominant antigens (e.g., major outer membrane protein), conditional GrgA-deficient, “maturation”-defective chlamydiae are potential candidates for life attenuated *Chlamydia* vaccines, provided that strategies are in place to fully prevent EBs from escaping the gene expression regulatory system in DOPE plasmid. To the minimum, the maturation-defective chlamydiae will serve as useful system for studying the roles of RBs in antichlamydial immunity.

## Materials and Methods

### Vectors

pTRL2-grgA-67m (Fig. S1), which carried a *grgA* allele with resistance to intron insertion between nucleotides 67 and 68, was constructed by assembling 3 DNA fragments using the NEBuilder HiFi DNA assembly kit (New England Biolabs). All 3 fragments were amplified from pTRL2-His-GrgA ^36^ using Q5 DNA polymerase (New England Biolabs). Fragment 1 was generated using primers pgp3-pgp4-F and His-RBS-R (Table S1). Fragment 2 was generated using primers RBS-His-F and GrgA-67-R (Table S1). Fragment 3 was generated using primers GrgA-67-F and pgp4-pgp3-R (Table S1). pDFTT3(aadA), a Targetron vector for disrupting chlamydial genes through group II intron insertional mutagenesis ^37^, was a generous gift from Dr. Derek Fisher (Southern Illinois University, IL). To construct pDFTT3(aadA)-GrgA-67 (Fig. S2), which was designed for disrupting the open reading frame of *grgA*, two PCR fragments were first generated using pDFTT3(aadA) as the template. Fragment 1 was obtained using primers GrgA67_IBS1/2 and the University primers (Table S1), while fragment 2 was obtained using primers GrgA67_EBS2 and GrgA67_EBS1/delta (Table S1). The two fragments were combined and subject to PCR extension. The resulting full-length intron-targeting fragment was digested with *Hind*III and *BsrG*I and subject to ligation with *Hind*III- and *BsrG*I-digested pDFTT3(aadA). The ligation product was transformed into *E. coli* DH5α, which was plated onto LB agar plates containing 500 µg/ml spectinomycin and 25 µg chloramphenicol. Authenticity of the insert in pDFTT3(aadA)-grgA-67m was confirmed using Sanger sequencing.

### Host cells and culture conditions

Mouse fibroblast L929 cells were used as the host cells for *C. trachomatis* transformation and preparation of EBs. Human vaginal carcinoma HeLa cells were used for experiments determining the effects of GrgA depletion on chlamydial growth and development. Both L929 and HeLa cell lines were maintained as monolayer cultures using Dulbecco’s modified Eagle’s medium (DMEM) (Sigma Millipore) containing 5% and 10% fetal bovine serum (vol/vol), respectively. Gentamicin (final concentration: 20 µg/mL) was used for maintenance of uninfected cells and was replaced with penicillin (10 units/mL) and/or spectinomycin (500 µg/mL) as detailed below. 37 °C, 5% CO2 incubators were used for culture uninfected and infected cells.

### Chlamydiae

Wildtype *C. trachomatis* L2 434/BU (L2) was purchased from ATCC. L2/cg-peig was derived by transforming L2 EBs with pTRL2-grgA-67m using calcium phosphate as previously described^36^. The transformation was inoculated onto L929 monolayer cells and selected with penicillin as previously described ^36^. L2/cgad-peig was derived by transforming L2/cg-peig with pDFTT3(aadA)-grgA-67m in the same manner. ATC was added to the cultures immediately after transformation to induce the expression of GrgA from pDFTT3(aadA)-grgA-67m. 12 hours later, spectinomycin D (final concentration: 500 µg/ml) was added to the culture medium to initiate selection^36^. L2/cgad-peig EBs were amplified using L929 cells and purified with ultracentrifugation through MD76 density gradients ^38^. Purified EBs were resuspended in sucrose-phosphate-glutamate (SPG) buffer; small aliquots were made and stored at -80 °C. Unless indicated otherwise, we added cycloheximide to all chlamydial cultures (final cycloheximide concentration in media: 1 µg/mL) to optimize chlamydia growth.

### Immunofluorescence staining

Near-confluent HeLa monolayers grown on 6-well plates were inoculated with L2/cgad-peig at MOI of 0.3 inclusion-forming units. Following 20 min centrifugation at 900 g, cells were cultured at 37 °C in media containing either 0 or 1 nM ATC for 30 h. The infected cells were then fixed with cold methanol, blocked with 10% fetal bovine serum prepared in phosphate-buffered saline (PBS), and stained successively with the monoclonal L21-5 anti-major outer membrane protein antibody^39,40^ and an FITC-conjugated rabbit anti-mouse antibody cells. Immunostained cells were finally counter-stained with 0.01% Evans blue (in PBS) before imaging under an Olympus IX51 fluorescence microscope. Red and green fluorescence images were acquired on an Olympus IX51 fluorescence microscope using a constant exposure time for each channel. Image overlay was performed using the PictureFrame software. The Java-based ImageJ software was then used to process the image^36^.

### IFU assays

L2/cgad-peig EB stock or frozen harvests of L2/cgad-peig cultured with or without ATC were thawed, 1-to-10 serially diluted, and inoculated onto L929 monolayers in medium containing 1 nM ATC and 1 µg/mL cycloheximide on a 96-w plates. Following 20 min centrifugation at 900 g, cells were cultured at 37 °C for 30 h. Cell fixation and antibody reactions were performed as described above. Immunostained inclusions were counted under the fluorescence microscope without Evan blue counter staining.

### Diagnostic PCR and DNA sequencing

For confirming and sequencing *grgA* alleles in the chromosome and plasmid, total DNA was extracted from ~ 1000 infected cells using the Quick-gDNA MiniPrep kit (Sigma Millipore) following manufacturer’s instructions. The resulting DNA was used for PCR amplification using the Taq DNA polymerase. DNA fragments resolved with electrophoresis of 1.2% Agarose gel were exercised and purified using the Gel Extraction kit (Qiagen) and subject to Sanger sequencing service provided by the Psomagen Service Center (New York).

### Quantification of genome copy numbers

To quantify genome copy numbers in cultures, infected cells on 6-, 12-, or 24-well plates were detached from the plastic using Cell Lifters (Corning). Cells and media were collected into Eppendorf tubes, centrifuged at 20,000 g at 4 °C. The supernatant was carefully aspirated. 100 µL alkaline lysis buffer (100 mM NaOH and 0.2 mM EDTA) was added into each tube to dissolve the cell pellets. Tubes were heated at 95 °C for 15 min and then placed on ice. 350 µL of H2O and 50 µl of 200 mM Tri-HCl (pH 7.2) were added into each tube and mixed. The neutralized extracts were used for qPCR analysis directly (1 µL/reaction) or after dilution with H2O.

### Western blotting

Detection of MOMP and GrgA was performed as previously described^36^. L929 cells grown on 6-well plates were infected with L2/cgad-peig and cultured with medium containing 1 nM ATC. At 15 h, 15 h 20 min, 15 h 40 min postinoculation, cells in selected wells were switched to ATC-free medium after 3 washes. At 16 hpi, cells in each well were harvested in 200 μL of 1X SDS-PAGE sample buffer, heated at 95 °C for 5 min, and sonicated for 1 min (5-s on/5-s off) at 35% amplitude. Proteins were resolved in 10% SDS-PAGE gels and thereafter transferred onto PVDF membranes. The membrane was propped with the monoclonal mouse anti-MOMP MC22 antibody^41^, stripped and reprobed with a polyclonal mouse anti-GrgA antibody^36^.

### Transmission electron microscopy

To visualize intracellular chlamydiae up to 36 hpi, L929 cell monolayers grown on 6-well plates were infected as described above and cultured with medium supplemented with or without 1 nM ATC. For cultures up to 36 h, cells were removed from the plastic surface using trypsin, collected in PBS containing 10% fetal bovine serum, and centrifuged for 10 min at 500 g. Pelleted cells were resuspended in EM fixation buffer (2.5% glutaraldehyde, 4% paraformaldehyde, 0.1 M cacodylate buffer) at RT, allowed to incubate for 2 h, and stored at 4 °C overnight. To visualize intracellular chlamydiae at 45 and 60 hpi, the above procedures resulted in lysis of infected cells and inclusions. To overcome this problem, cells grown on glass coverslips were infected with and fixed without trypsinization. To prepare samples for imaging, cells were first rinsed in 0.1 M cacodylate buffer, dehydrated in a graded series of ethanol, and then embedded in Eponate 812 resin at 68 °C overnight. 90 nm thin sections were cut on a Leica UC6 microtome and picked up on a copper grid. Grids were stained with Uranyl acetate followed by Lead Citrate. TIFF images were acquired on a Philips CM12 electron microscope at 80 kV using an AMT XR111 digital camera. RB diameters were measured using ImageJ software [46].

## ACKNOWLEDGEMENTS

We thank Rajesh Patel for processing electron microscopy samples, Dr. Derek Fisher (Southern Illinois University) for pDFTT3(aadA) and Drs. Harlan D. Caldwell (National Institute of Allergy and Infectious Diseases) and Guangming Zhong (University of Texas Health Sciences Center at San Antonio) for antibodies. We also acknowledge Tara Kehair’s participation in the analysis of electron microscopic images. This work was supported by grants from the National Institutes of Health (Grant # AI AI140167 and AI 154305 to HF).

